# Familial influences on Neuroticism and Education in the UK Biobank

**DOI:** 10.1101/582627

**Authors:** R. Cheesman, J. Coleman, C. Rayner, K.L. Purves, G. Morneau-Vaillancourt, K. Glanville, S.W. Choi, G. Breen, T.C. Eley

## Abstract

Genome-wide studies often exclude family members, even though they are a valuable source of information. We identified parent-offspring pairs, siblings and couples in the UK Biobank and implemented a family-based DNA-derived heritability method to capture additional genetic effects and multiple sources of environmental influence on neuroticism and years of education. Compared to estimates from unrelated individuals, heritability increased from 10% to 27% and from 19% to 57% for neuroticism and education respectively by including family-based genetic effects. We detected no family environmental influences on neuroticism, but years of education was substantially influenced by couple similarity (38%). Overall, our genetic and environmental estimates closely replicate previous findings from an independent sample, but more research is required to dissect contributions to the additional heritability, particularly rare and structural genetic effects and residual environmental confounding. The latter is especially relevant for years of education, a highly socially-contingent variable, for which our heritability estimate is at the upper end of twin estimates in the literature. Family-based genetic effects narrow the gap between twin and DNA-based heritability methods, and could be harnessed to improve polygenic prediction.

## Introduction

Heritability measures the proportion of individual differences in a trait explained by genetic variation, and is traditionally estimated by comparing the resemblance of identical and non-identical twins (Knopik et al. 2017). Researchers can also now estimate single nucleotide polymorphism (SNP) heritability, the variance explained by the additive effects of common genetic variants tagged by a genotyping array (Yang et al. 2010; Yang et al. 2011). SNP heritability is expected to be less than twin heritability, since the former only estimates the additive effects of measured common variants, plus variants that are correlated (i.e. in linkage disequilibrium) with them, rather than the influence of DNA sequence differences that are rare and/or not well tagged by genotyping arrays. Since genome-wide association studies (GWAS) also only consider the additive effects of common variants, SNP heritability provides an estimate of the total genetic effect that could be identified with well-powered association studies of a given phenotype in a given population. Given the importance of SNP heritability, researchers have investigated approaches to maximise the accuracy of estimates, beyond increasing sample sizes or denser genotyping (van den Berg et al. 2014; Laurin et al. 2015; Cheesman et al. 2018; van der Sluis et al. 2010).

The dominant method for the estimation of SNP heritability, Genomic-RElatedness-based restricted Maximum-Likelihood (GREML), takes advantage of small genetic differences among many unrelated individuals to predict trait similarity (Yang et al. 2011). The derived SNP heritability estimates reflect genetic effects tagged by common SNPs. The effects of genetic variants that are not in linkage disequilibrium with common genotyped SNPs will not be captured using this method. However, when GREML is applied to family data, the higher genetic relatedness among relatives increases the correlation between genotyped SNPs and causal variants, because they are more likely to be inherited together (Zaitlen et al. 2013; Xia et al. 2016). This increase in linked variants helps to capture additional genetic variation not normally picked up in population studies of unrelated people, such as rare variants, copy number variants, and structural variants.

An extension of the method that uses family data, GREML-KIN, allows the estimation of two categories of genetic influence: population-level common variant heritability, plus additional heritability that is associated with kinship (Xia et al. 2016; Zaitlen et al. 2013). The first estimate is similar to that derived from GREML using unrelated individuals. The latter heritability estimate captures additional family-based effects, due to the increased correlation between genotyped SNPs and causal variants among relatives. The GREML-KIN method also allows for effects of environment sharing amongst family members, siblings and couples. This inclusion is important as it attempts to remove confounding that results from people who are more genetically related having more similar environments and higher phenotypic resemblance than people who are less related.

One study applied the GREML-KIN method to neuroticism and years of education (Hill et al. 2018) in Generation Scotland, a large family-based study (Smith et al. 2006). Their sample was selected to capture dense kinships, where many individuals have siblings, parents and spouses who are also study participants. Neuroticism is a personality trait characterised by readily experiencing negative emotions. It is a strong predictor of common mental disorders including depression, anxiety, schizophrenia and substance abuse, and is associated with important life outcomes such as occupational attainment, divorce, and mortality (Ormel et al. 2013). Years of education is also a complex trait that shows significant associations with diverse social, economic and health outcomes (Mackenbach et al. 2008). Twin studies have repeatedly demonstrated substantial heritability for neuroticism (40-60%) (Hettema et al. 2006) and educational attainment (40-50%) (Branigan et al. 2013). These are both broad phenotypes that index a wide range of important traits, and are available in numerous very large datasets. As a result, they have been used to perform some of the largest GWA studies of psychological traits (N=449,484 for neuroticism (Nagel et al. 2018; Luciano et al. 2018); N=1.1 million for years of education (Lee et al. 2018)). Nonetheless, common SNPs only explain ∼10% of the variance in neuroticism (Luciano et al. 2018; Nagel et al. 2018). Estimates of SNP heritability for years of education also fall substantially short of twin estimates (14.7%; Lee et al. 2018).

GREML-KIN analyses in Generation Scotland revealed a large increase in heritability compared to a standard GREML analysis of unrelated individuals (Hill et al. 2018). For neuroticism, the total heritability from the best-fitting model was 30%, primarily accounted for by kin-based genetic effects (19%), as well as common variant effects tagged in studies of unrelated individuals (11% - akin to SNP heritability). They detected no family environment effects. For years of education, there was a strong kin genetic component (28%) in addition to common genetic influence (16%), plus substantial ‘environmental’ effects of sibling and couple similarity (11% and 31%, respectively). The findings align well with evidence from twin studies that the family environment influences education-related outcomes (∼36% of the variance; Branigan et al. 2013) but not neuroticism (Polderman et al. 2015). If these results are replicated, then the total DNA-based heritability of neuroticism and education would be close to twin study estimates. Of note, for each, the larger component of the genetic contributions results from less common variants not identified in genomic studies of unrelated individuals. Moreover, a replication of these results would suggest that most of the variance in education (86%) can be captured with measured parameters. Notably, the authors also found that rarer variants (0.1%-1% in frequency) explained 12% of the variance in education, but did not influence variation in neuroticism (Hill et al. 2018).

Our study aimed to estimate familial influences on neuroticism and years of education in the UK Biobank, using GREML-KIN. We utilise the presence of thousands of family members in the UK Biobank to shed light on the genetic and environmental architecture of these two phenotypes. To robustly replicate the previous Generation Scotland study, we ensured that phenotype definitions were as similar as possible, and that there was no sample overlap. Based on previous research, we hypothesised that neuroticism and years of education would show increased heritability by exploiting the higher linkage disequilibrium within families. Our secondary analyses aimed to validate our kin-based estimates and specifically quantify the contribution of rarer genetic variants to the heritability of neuroticism and years of education. For this, we used the GREML-MS method, which stratifies imputed genetic variants by their frequency to allow estimation of the variance explained by rarer variants (Yang et al. 2015).

## Methods

### Sample

Analyses were conducted using the UK Biobank, a resource containing rich phenotype and genotype data on ∼500,000 individuals aged between 40 and 70 (Allen et al. 2014). Analyses were restricted to white British individuals. We identified families and restricted heritability analyses to individuals with at least one family member in the UK Biobank, as well as phenotype data on neuroticism and/or years of education. Previous publications suggested a sample size of ∼40,000 pairs of family members (parent-offspring, siblings, and couples) within the full dataset (Bycroft et al. 2018).

### Genotyping

Genome-wide genetic data from the full release of the UK Biobank data were collected and processed according to the quality control pipeline (Bycroft et al. 2018). For primary GREML-KIN analyses, we used genotyped or imputed SNPs with minor allele frequency >0.01 and imputation confidence (INFO) score >0.4, indicating well imputed variants. Due to computing memory constraints, we used PLINK2 to prune down to 241678 variants in approximate linkage equilibrium using an r^2^ threshold of 0.2 (Chang et al. 2015) before calculating genetic relatedness matrices.

### Measures

Neuroticism was measured as a continuous trait, captured with 12 questionnaire items such as “Does your mood often go up and down?”, “Would you call yourself tense or ‘highly strung’?”. This trait was defined previously in the UK Biobank (Smith et al. 2016; Smith et al. 2013).

The years of education variable was defined according to ISCED categories, as in previous genomic studies in the UK Biobank and other samples (Hill et al. 2018; Lee et al. 2018). The six response categories were: none of the above (no qualifications) = 7 years of education; CSEs or equivalent = 10 years; O levels/GCSEs or equivalent = 10 years; A levels/AS levels or equivalent = 13 years; other professional qualification = 15 years; NVQ or HNC or equivalent = 19 years; college or university degree = 20 years of education. To test whether the number of response categories affected heritability estimates, as has been shown previously in the UK Biobank (Lee et al. 2018), we ran sensitivity analyses using a ‘coarsened’ years of education variable, plus a binary variable reflecting college completion (see Supplementary Figure 1).

**Figure 1:**
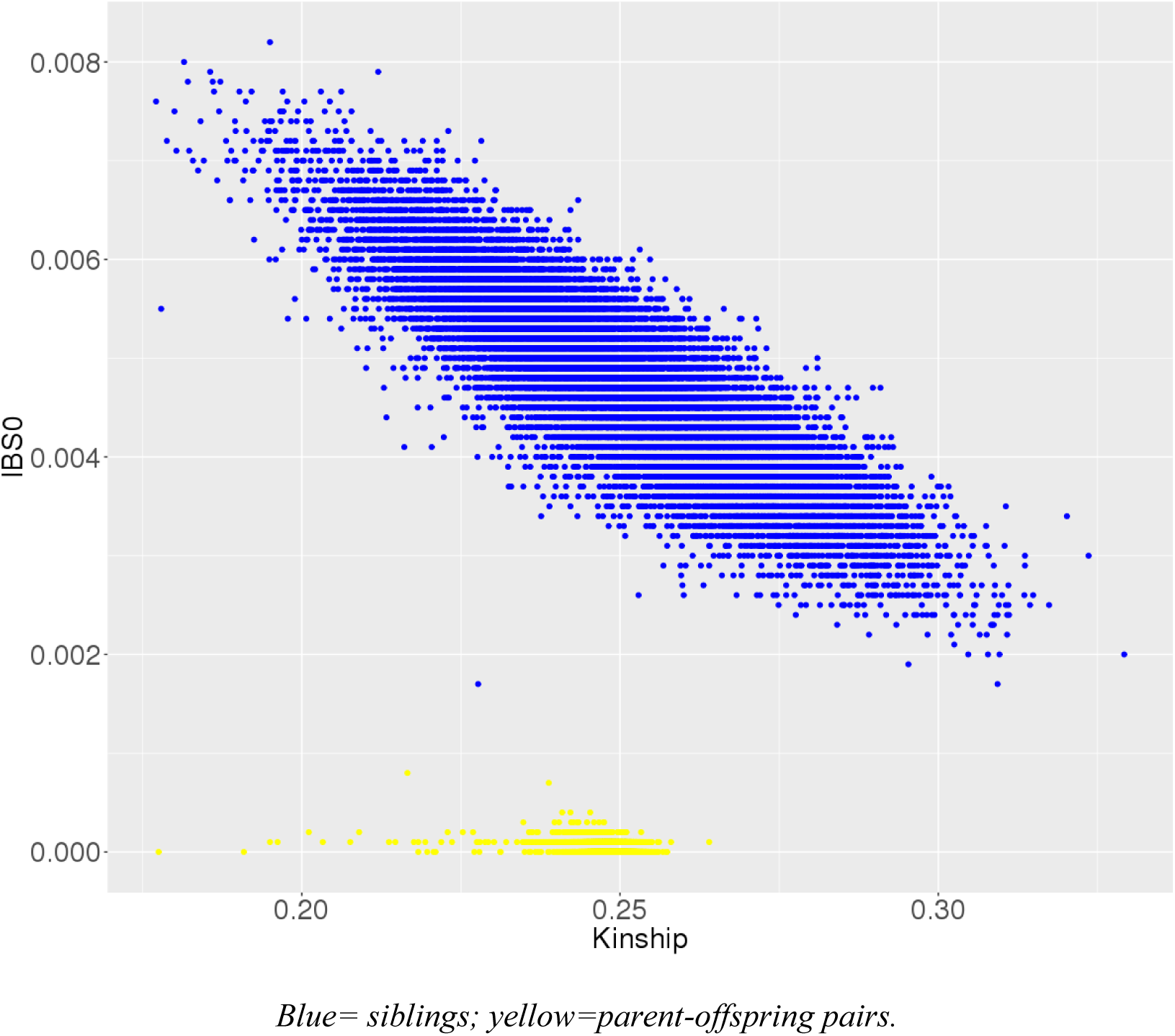
Kinship plotted against IBS0 for all first-degree relatives in the UK Biobank. Blue= siblings; yellow=parent-offspring pairs.

We regressed out the effects of age, sex and population stratification (using 20 principal components of the genetic relatedness matrix from GCTA for the latter) prior to model fitting.

### Analyses

#### Identification of family members

Sibling and parent-offspring pairs were identified using relatedness files (KING n.d.) received with the UK Biobank data. Relatedness between two individuals is summarised by a kinship coefficient, which is defined as the probability that a random allele from an individual is identical by descent (IBD) with an allele at the same locus from the other individual (i.e. identical and inherited from a common ancestor). For example, in parent-offspring duos, kinship is ∼0.25, as it is the probability that a random allele in a child is from one specific parent (0.5 since humans are diploid) multiplied by the probability that the parental allele from that parent is passed to the child (0.5; independent to the first probability). To allow for normal variation in within-pair similarity, first-degree relatives are therefore defined as pairs that have a pairwise kinship coefficient of >= 0.177 and <= 0.354.

To distinguish parent-offspring pairs from sibling pairs, we plotted the proportion of SNPs with *zero* identity-by-state (IBS0) within the kinship bounds of 0.177-0.354 (Figure 1). IBS describes the probability that alleles are the same regardless of common ancestry. When comparing two individuals, variants are termed *IBS0* if neither allele is shared by the pair. Parent-offspring pairs have IBS0=∼0 since they share one allele inherited by descent (IBD) in all positions on autosomes. In other words, they are unlikely to share *zero* variants. In contrast, siblings have a higher pairwise IBS0.

Couples were identified as pairs of unrelated opposite-sex individuals matching exactly on a string of household variables: social deprivation (Townsend Deprivation Index), assessment centre, income, time at address, smoker in household, type of accommodation, relatives in household, number in household. This approach of matching on household variables was used in a recent study of assortative mating in the UK Biobank (Yengo et al. 2018). We note that there is potential for type 1 error: it is possible, especially in densely populated areas, that people could match on all eight variables by chance.

### Kin-based SNP heritability method accounting for environmental similarity: GREML-KIN

GREML requires the calculation of genetic similarity for each pair of individuals across genotyped variants. This matrix of genomic similarity is compared to a matrix of pairwise phenotypic similarity using a random-effects mixed linear model, such that the variance of a trait can be decomposed into genetic and residual components, using maximum likelihood. Ordinarily, GREML is applied in samples of unrelated individuals and has a single common genetic variance component.

GREML-KIN is an extension of GREML that estimates the variance explained by multiple genetic and non-genetic sources. The method uses a linear mixed model to fit five matrices: G= common genotyped SNP effects; K= kin genetic effects; F= nuclear family (siblings, parent-offspring, and couples); S= siblings; C= couples. For the G matrix, we calculated genetic similarity for all possible pairs of individuals across all genotyped SNPs. As GREML-KIN allows for effects of the family environment, no relatedness cut-off was applied to the G matrix (unlike the standard GREML model applied only to unrelated individuals, where a cut-off of <0.025 is typically used). The K matrix is simply a modified G matrix, containing only information on relatives (cut-off >0.025), since values for unrelated pairs are set to 0. Family, sibling and couple (F, S and C) similarity matrices were created in the format required for GCTA. Elements in the genomic relatedness matrix were replaced by 0 if a pair did not have the specific relationship; and 1 if a pair do have the relationship, or for elements representing individuals’ relatedness to themselves.

Importantly, the matrices are not purely ‘genetic’ and ‘environmental’. The sibling and couple environment sharing matrices likely pick up variance due to other processes that inflate covariance between relatives, including dominance and assortative mating, which refers to greater similarity between partners than is expected by chance. This can result from multiple mechanisms, including direct choice, social homogamy, and/or convergence over time due to shared environments.

The genetic variance components are likely to include some bias from the indirect effects of genetic variants shared with relatives, which are environmental from the perspective of the individual under study (see Discussion for a detailed consideration of confounding due to gene-environment correlation). The residual component includes any other source of variance that are not captured by the G, K, F, S or C matrices. The residual variance includes other environmental influences (especially idiosyncratic, individual-specific environments or perceptions that are not shared by family members) and error.

To identify the best-fitting model for each trait, we ran a model for every possible combination of variance components (31 models), and compared them with backwards stepwise likelihood ratio testing, starting with the full model and dropping non-significant parameters.

We compared GREML-KIN results to a standard GREML model in a subset of unrelated individuals from the family-based analyses. The standard GREML model uses a single genomic relatedness matrix with a cut-off to exclude one from each pair of related individuals (cutoff >0.025). This approach therefore only detects population-level additive genetic effects tagged by common genotyped SNPs, plus potential confounding from indirect genetic effects and population stratification. The residual component contains other sources of variance: gene–environment interaction, error, plus all of the environmental influence, rare variant effects that are not captured when using an unrelated population sample, and non-additive genetic effects.

### GREML-MS to investigate the effects of less common variants

In our GREML-MS analyses, we took imputed whole genome sequence data and ran quality control to include variants across the allele frequency spectrum that were imputed with high confidence (INFO score >0.80). Multiallelic variants were removed. We created six genome-wide genetic relatedness matrices, one for each allele frequency bin (non-overlapping). The bins/GRMs contained variants with minor allele frequency ranges of: 0.001-0.01, 0.01-0.1, 0.1-0.2, 0.2-0.3, 0.3-0.4, 0.4-0.5, respectively. These thresholds match the previous study in Generation Scotland. All matrices included the same number of unrelated individuals with phenotype data and with at least one family connection as in standard GREML. The matrices are simultaneously fitted using a linear mixed model.

### Sample independence and software

To ensure that our results were independent from the previous Generation Scotland study, we compared checksums for both samples to identify and remove overlapping participants. A checksum is the sum of nine numbers taken from binary genotype files. Checksums were obtained from Generation Scotland without accessing genotype data directly. We then ran checksums in the UK Biobank (after ensuring quality control of genomic data was the same), using a script from the Broad Institute, which is available online: https://personal.broadinstitute.org/sripke/share_links/checksums_download/outdated_readme/id_geno_checksum.v2.

We used the following software in our analyses: identification of family members was performed using R; construction of genomic relationship matrices was done in GCTA; family, sibling and couple similarity matrices were made in bash; GREML analyses were conducted in GCTA. Scripts are available from the lead author on request. The UK Biobank is a public dataset available to all bona fide researchers (with funds to pay the access fee).

## Results

### Identification of family members

Columns 2-4 of Table 1 (bold) show how many pairs of the three types of family members we identified with available data on neuroticism and years of education. The numbers of family relationships closely matched findings from previous publications (Yengo et al. 2018; Bycroft et al. 2018). Column 5 contains the sample sizes of family pairs (the total of couple, sibling and parent-offspring pairs). From the number of family pairs, we derived the number of families (column 6), or in other words, the number of discrete sets of individuals who have at least one connection. The number of unique individuals (column 7) represents the total number of participants with at least one family member, after removing double-counted individuals who have multiple family members in the sample.

**Table 1:** Sample sizes for different family relationships in the UK Biobank.

| *Phenotype* | *Couple (pairs)* | *Sib (pairs)* | *Parent-offspring*<br>*(pairs)* | *Nuclear (pairs)* | *Families* | *Unique individuals* |
| --- | --- | --- | --- | --- | --- | --- |
| Neuroticism | **16454** | **14880** | **4121** | 35455 | 31783 | 66118 |
| Education | **23205** | **21568** | **5915** | 50688 | 45160 | 93150 |

The discrepancies between the numbers of individuals, families and nuclear pairs reflects that most people only have one family member in the study. This contrasts to samples with dense kinship networks such as Generation Scotland, where many individuals have siblings, parents and spouses who are also study participants. As described in the Methods, we distinguished parent-offspring and sibling pairs according to their IBS0 (Figure 1). We chose a threshold of IBS0>0.001 to define siblings (blue) separately from parent-offspring pairs (yellow).

Phenotypic correlations for neuroticism were 0.03 for couples, 0.14 for siblings, and 0.13 for parent-offspring pairs. Phenotypic correlations for years of education were 0.38 for couples, 0.30 for siblings, and 0.26 for parent-offspring pairs.

### Kin-based SNP heritability method accounting for environmental similarity: GREML-KIN

Figure 2 shows results for the full GREML-KIN models for neuroticism and education years. For neuroticism, the full model indicated that the variance explained by common genetic and kin-based variants is 11% (se=0.01) and 18% (se=0.08), respectively (29% in total, first two bars). Bars 3-5 demonstrate that there is no significant influence of family, sibling or couple similarity. For neuroticism, the selected model gives similar results and contains common SNP and kin-based genetic influences (11% (se=0.01) and 16% (0.02) respectively). Compared to this best-fitting model, the inclusion of matrices to control for the influence of family environments (in the full model) does not reduce heritability. The total heritability of neuroticism when including relatives (27%) is substantially higher than in our analysis of unrelated individuals: 10% (se=0.01; N= 44694 unrelated individuals; Supplementary Table 1).

**Figure 2:**
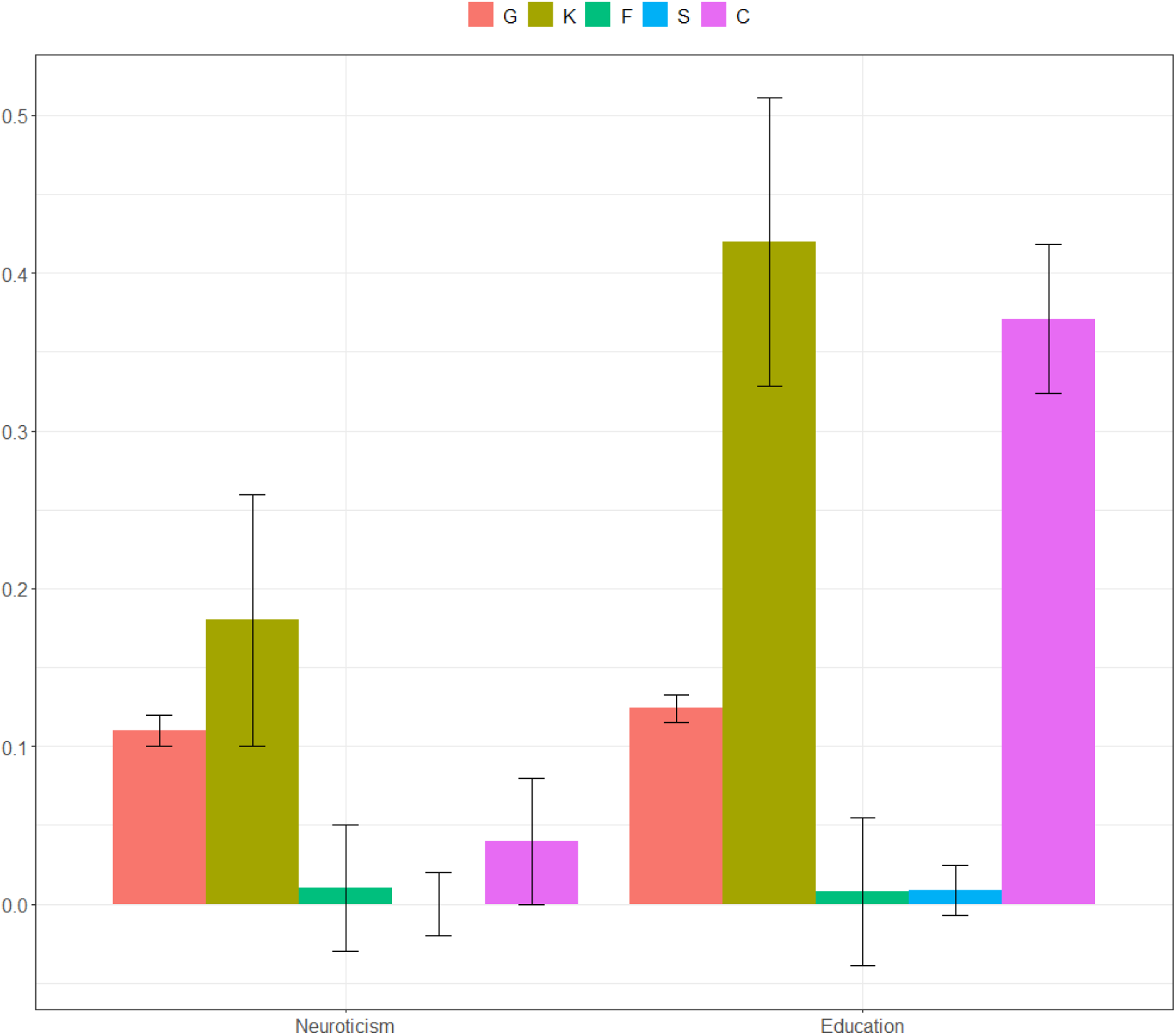
variance component estimates for neuroticism and education, plus standard errors. G= population-level effects of common genotyped SNPs; K= kin-based genetic effects; F= effects of nuclear family (siblings, parent-offspring, couple) similarity; S= effects of sibling similarity; C= effects of couple similarity. Note that for neuroticism, the estimates from our full model for the F, S, and C components are all non-significant, and standard errors cross zero. For education, the F and S components are non-significant.

Figure 2 also shows the results for years of education from our full model. Due to computational memory limitations, it was necessary to run models for education in two parts (explained below). We report results for the first part in the main text, and the smaller second part in Supplementary Table 1. The heritability of years of education in our selected model reached 57%, made up of 12% common SNPs at the population level (se=0.01) and 45% kin-based effects (se=0.02). This increases heritability considerably from 19% (se=0.01) in unrelated individuals. The final model for education also contains a couple similarity effect of 38% (se=0.01) in addition to the common and kin-based genetic influences.

See Supplementary Table 1 for full model-fitting results, including all sub-models and fit statistics. As noted above, we ran two sets of GREML-KIN models in mutually exclusive samples to reduce the computational burden. The first used the same matrices as in analyses of neuroticism (N= 66118). The second used new matrices including individuals who have education data and at least one family member, and who were not included in the neuroticism matrices. In defining these groups, we ensured that individuals in the same family were kept together, which led to sample sizes of 62353 and 30797 for the two groups, respectively. Results for the second part were very similar to those from the first, with estimates from the best-fitting model (GKC), at 12% (se=0.02), 49% (0.03) and 37% (0.01), respectively (Supplementary Table 1).

Supplementary Figure 1 gives GREML-KIN results for alternative education phenotypes (years of education with fewer categories, and degree/college completion). Estimates differed only slightly, and the conclusions remain the same.

### GREML-MS to investigate the effects of less common variants

Table 2 shows our estimates of the contribution of variants of different allele frequencies to neuroticism and years of education. Results suggest genetic influences across the allele frequency spectrum for both traits. For neuroticism, there is no evidence of any contribution made by the SNPs in our lowest minor allele frequency (MAF) bin (0.001–0.01), and all variants only explained 10% (se=0.01) of phenotypic variation. For education, variants tagged by low frequency SNPs (MAF between 0.001–0.01, and 0.01– 0.1) make a modest contribution to phenotypic variation, and all variants explained 22% of the phenotypic variance.

**Table 2:** Results of GREML-MS variance components analyses for neuroticism and education using six minor allele frequency bins.

|  | MAF | 0.001–0.01 | >0.01–0.1 | >0.1–0.02 | >0.2–0.3 | >0.3–0.4 | >0.4–0.5 | Total |
| --- | --- | --- | --- | --- | --- | --- | --- | --- |
|  | Number of<br>SNPs | 3,544,806 | 3,512,584 | 1,469,557 | 1,045,242 | 885,812 | 809,397 | 11,267,398 |
| Neuroticism | h<sup>2</sup> | 0.00 (0.01) | 0.02 (0.01) | 0.01 (0.01) | 0.03 (0.01) | 0.03 (0.01) | 0.02 (0.004) | 0.10 (0.01) |
| Education | h<sup>2</sup> | 0.06 (0.01) | 0.04 (0.01) | 0.03 (0.01) | 0.03 (0.01) | 0.03 (0.01) | 0.03 (0.01) | 0.22 (0.02) |

MAF= minor allele frequency, h<sup>2</sup> (se) = variance explained by SNPs in MAF bin, plus standard error.

Checksum analyses indicated that there were no family members with phenotype data in the UK Biobank who were also in Generation Scotland. We can therefore be confident that our results are independent of the previous study (Hill et al. 2018).

## Discussion

In this study, we capitalised on the family information and genome-wide genotypes in the UK Biobank to estimate genetic and family environmental influences on two psychologically, socially and economically important quantitative phenotypes: neuroticism and years of education. Inclusion of family-based genetic effects substantially increases the heritabilities of neuroticism and education from 10% and 19% in standard analyses of unrelated individuals, to 27% and 57% respectively. Our estimates closely match previous findings from an independent sample (Generation Scotland; Hill et al. 2018), and bring the variance explained by measured DNA alone closer to twin estimates (40-60% and 40-50% for neuroticism and education, respectively). The additional family-based influences likely include rare variants, copy number variants, and structural variants.

Turning to our non-genetic findings, we did not detect any influence of nuclear family, sibling or couple similarity on variation in neuroticism. This is consistent with evidence from twin studies against shared environmental contributions to personality (Nivard et al. 2015; Hettema et al. 2006), and with the previous study using the same method (Hill et al. 2018). For education years, we found an effect of couple similarity (38%), but not family or sibling similarity. This couple effect likely captures assortative mating (the education years is likely to have occurred before the couple became a couple) and family environment effects on education. This aligns with previous evidence for assortative mating for education in the UK Biobank (Yengo et al. 2018; Robinson et al. 2017) and other samples (Domingue et al. 2014).

The main limitation of our approach is its potential inability to distinguish genetic from environmental influences. There are two key reasons why our heritability estimates could contain residual environmental confounding: lack of dense relationships in the sample, and gene-environment correlation. Firstly, despite containing large numbers of different types of relatives, the number of relationships per individual in the UK Biobank is lower than in a purpose-built kinship sample like Generation Scotland. This likely results in lower power in the UK Biobank to detect influences of family similarity, especially if they are small in magnitude, and reduces power to separate confounding factors. Recent inter-generational structural equation modelling research has shown that statistical power to distinguish variance components increases exponentially with the addition of siblings to analyses – more so than including additional families in analyses (McAdams et al. 2018). We hope that these demands will be met with the development of intergenerational population samples with large amounts of genomic family data.

Secondly, residual environmental confounding is likely to remain in our GREML-KIN heritability estimates due to gene-environment correlation. Gene-environment correlation among parents and offspring reflects that genetic variants in the parents do not only have direct effects on offspring traits by being transmitted, but they also have indirect influences on offspring traits through the environment that they provide for their children. Simulations suggest that although GREML-KIN gives unbiased heritability estimates when family environmental influences are present, it over-estimates heritability if phenotypes are also substantially influenced by the indirect effects of transmitted alleles, also known as passive gene-environment correlation (Young et al. 2018). Methods exist to separate the direct heritability (17% for educational attainment in Young et al. (2018)) from indirect genetic effects, but we are underpowered in the UK Biobank to apply these approaches, which currently rely on very large numbers of siblings or parent-offspring trios (Eaves et al. 2014; Visscher et al. 2006; Young et al. 2018). Intergenerational and/or sibling data should be used to distinguish direct from indirect influences, if we are to understand causal mechanisms in the aetiology of neuroticism and education and other complex, socially-contingent traits, and to estimate accurately the potential for ‘environmental’ modification. In addition, one study found that controlling for urban childhood residence, a proxy for a range of environmental factors, reduced the SNP heritability of education (but not the heritabilities of height or body mass index) above and beyond other controls for population stratification (Conley et al. 2014). Particularly for education, geographic population differences, which are interlinked with indirect effects from relatives, complicate our interpretations of heritability estimates.

Although these potential sources of environmental confounding indicate that our interpretations should be cautious, our heritability estimates are notably similar to those from twin studies. Twin heritability estimates are viewed as the gold standard because they capture the effects of any DNA sequence differences, and because they are well-replicated across decades of studies (Polderman et al. 2015; Plomin et al. 2016). Importantly, twin heritability estimates are not entangled with passive gene-environment correlation, which instead loads as shared environment. It therefore appears likely that our family based genomic heritability estimates are capturing some genetic variation that is missed in studies of unrelated individuals, and that the slightly increased heritability for education in this study compared to twin-based estimates in the literature (57% vs 40-50%, respectively) is due to passive gene-environment correlation.

This leads us to consider the extent to which the large contribution of family-associated genetic influence is explained by rarer variants. For education, some additional lower frequency effects are captured by variants in our allele frequency partitioned genomic relatedness matrices. For Neuroticism, the same GREML-MS heritability analyses indicated less rare genetic influence than expected based on the large kin-genetic component from the GREML-KIN model-fitting. The lack of agreement between GREML-MS and GREML-KIN heritability estimates might be because the former method is less able to capture very rare effects (allele frequency <0.001). Moreover, it has been shown that GREML-MS analysis of imputed rather than whole genome sequence data underestimates heritability when a trait is strongly influenced by rare variants (Evans, Tahmasbi, Vrieze, et al. 2018). For neuroticism, the gain in heritability from GREML-KIN despite the lack of variance in the 0.001-0.01 frequency bin is likely to be partly explained by very rare, non-additive, and structural variation.

Evidence from other studies suggests an important role for rare variants, which will be further elucidated with the increasing availability of whole genome sequence data. Multifactorial traits such as personality and educational attainment are thought to have complex genetic aetiologies, consisting of interplay between common and rare variation. Recent exome sequencing work has demonstrated rare coding variant effects on many psychiatric traits (that are closely related to neuroticism; (Ganna et al. 2018)). Other evidence for interactions between common variants across the genome and rare copy number variants in schizophrenia (Bergen et al. 2018) shows the importance of examining different types of genetic variation together. Importantly, twin and pedigree estimates of heritability can be recovered by using whole genome sequence data (rather than imputed data) in GREML analyses across the phenome, although much larger sample sizes are needed to detect rare genetic influences (Yang et al. 2015; Evans, Tahmasbi, Vrieze, et al. 2018).

The additional influences correlated with kinship detected in this study indicate a chance to improve prediction using DNA. Polygenic risk scores combine genome-wide genotype data into a single variable that measures genetic liability to a trait. They are generated by summing risk alleles carried by an individual, weighted by the effect size from the discovery GWAS (Lewis and Vassos 2017). The prediction accuracy of polygenic scores is a function of the SNP heritability of the target sample, and of the SNP heritability of, and genetic correlation with, the phenotype in the GWAS discovery sample (de Vlaming et al. 2017).

Polygenic scores for neuroticism currently explain maximum 4.2% of the phenotypic variance in independent samples, which is a fraction of the common SNP heritability (∼10%; Nagel et al, 2018). For years of education, polygenic scores explain more than 10% of the phenotypic variance, around two thirds of the SNP heritability of 15%. Increasing sample sizes and phenotypic homogeneity for GWAS of unrelated population samples will continue to narrow this gap between polygenic score prediction and SNP heritability. However, the population-level common variant association approach will be limited for educational attainment, given the relatively high variance already explained by polygenic scores. For both traits, genomic prediction could be improved by leveraging family information (Lee et al. 2017). Inclusion of relatives could help to capture additional effects of typically untagged variants that are likely rare and possibly even family-specific. Importantly, for the purpose of prediction, genetic predictors need not be, and already are not, ‘pure’ indices of individual genetic potential. Polygenic scores will be more predictive if they can better leverage indirect genetic effects from relatives to tag influential aspects of the environment, for example by combining parent and child polygenic scores within a single model.

In summary, we provide evidence for substantial family-based and common genetic effects on neuroticism and years of education in the UK Biobank. These results motivate the recruitment of samples with dense kinships and methods to leverage genomic family information, whilst understanding gene-environment interplay.

## Acknowledgments

We would like to thank the scientists involved in the construction of the UK Biobank and all of the participants who have shared their life experiences with investigators in the UK Biobank. This research has been conducted using the UK Biobank Resource, under the application 18177 (with thanks to Paul F. O’Reilly). This study represents independent research part funded by the National Institute for Health Research (NIHR) Biomedical Research Centre at South London and Maudsley NHS Foundation Trust and King’s College London. The views expressed are those of the author(s) and not necessarily those of the NHS, the NIHR or the Department of Health and Social Care. High performance computing facilities were funded with capital equipment grants from the GSTT Charity (TR130505) and Maudsley Charity (980). T.C. Eley is part funded by a program grant from the UK Medical Research Council (MR/M021475/1). R. Cheesman is supported by an ESRC studentship. C. Rayner is supported by a grant from Fondation Peters to T.C. Eley and G. Breen. K.L. Purves is part supported by a grant from the Alexander Von Humboldt Foundation and UK Medical Research Council (MR/M021475/1). G. Morneau-Vaillancourt is supported by a studentship from the Quebec Network on Suicide, Mood Disorders and Related Disorders. S.W. Choi. is funded from the UK Medical Research Council (MR/N015746/1). KG is supported by a PhD studentship awarded from the UK Medical Research Council. We thank David M. Howard for assistance with checksums for Generation Scotland, and Loic Yengo for sharing information on identification of couples.

The authors declare no conflicts of interest.

## References

Allen, N.E., Sudlow, C., Peakman, T., Collins, R. and UK Biobank 2014. UK biobank data: come and get it. Science Translational Medicine 6(224), p. 224ed4.

Benyamin, B., Visscher, P.M. and McRae, A.F. 2009. Family-based genome-wide association studies. Pharmacogenomics 10(2), pp. 181–190.

Bergen, S.E., Ploner, A., Howrigan, D., et al. 2018. Joint contributions of rare copy number variants and common snps to risk for schizophrenia. The American Journal of Psychiatry, p. appiajp201817040467.

Branigan, A.R., McCallum, K.J. and Freese, J. 2013. Variation in the Heritability of Educational Attainment: An International Meta-Analysis. Social Forces 92(1), pp. 109–140.

Bycroft, C., Freeman, C., Petkova, D., et al. 2018. The UK Biobank resource with deep phenotyping and genomic data. Nature 562(7726), pp. 203–209.

Chang, C.C., Chow, C.C., Tellier, L.C., Vattikuti, S., Purcell, S.M. and Lee, J.J. 2015. Second-generation PLINK: rising to the challenge of larger and richer datasets. GigaScience 4, p. 7.

Cheesman, R., Major Depressive Disorder Working Group of the Psychiatric Genomics Consortium, Purves, K.L., et al. 2018. Extracting stability increases the SNP heritability of emotional problems in young people. Translational psychiatry 8(1), p. 223.

Colodro-Conde, L., Rijsdijk, F., Tornero-Gómez, M.J., Sánchez-Romera, J.F. and Ordoñana, J.R. 2015. Equality in educational policy and the heritability of educational attainment. Plos One 10(11), p. e0143796.

Conley, D., Siegal, M.L., Domingue, B.W., Mullan Harris, K., McQueen, M.B. and Boardman, J.D. 2014. Testing the key assumption of heritability estimates based on genome-wide genetic relatedness. Journal of Human Genetics 59(6), pp. 342–345.

de Vlaming, R., Okbay, A., Rietveld, C.A., et al. 2017. Meta-GWAS Accuracy and Power (MetaGAP) Calculator Shows that Hiding Heritability Is Partially Due to Imperfect Genetic Correlations across Studies. PLoS Genetics 13(1), p. e1006495.

Domingue, B.W., Fletcher, J., Conley, D. and Boardman, J.D. 2014. Genetic and educational assortative mating among US adults. Proceedings of the National Academy of Sciences of the United States of America 111(22), pp. 7996–8000.

Eaves, L.J., Pourcain, B.S., Smith, G.D., York, T.P. and Evans, D.M. 2014. Resolving the effects of maternal and offspring genotype on dyadic outcomes in genome wide complex trait analysis (“M-GCTA”). Behavior Genetics 44(5), pp. 445–455.

Evans, L.M. and Keller, M.C. 2018. Using partitioned heritability methods to explore genetic architecture. Nature Reviews. Genetics 19(3), p. 185.

Evans, L.M., Tahmasbi, R., Jones, M., et al. 2018. Narrow-sense heritability estimation of complex traits using identity-by-descent information. Heredity 121(6), pp. 1–15.

Evans, L.M., Tahmasbi, R., Vrieze, S.I., et al. 2018. Comparison of methods that use whole genome data to estimate the heritability and genetic architecture of complex traits. Nature Genetics 50(5), pp. 737–745.

Ganna, A., Satterstrom, F.K., Zekavat, S.M., et al. 2018. Quantifying the Impact of Rare and Ultrarare Coding Variation across the Phenotypic Spectrum. American Journal of Human Genetics 102(6), pp. 1204–1211.

Hettema, J.M., Neale, M.C., Myers, J.M., Prescott, C.A. and Kendler, K.S. 2006. A populationbased twin study of the relationship between neuroticism and internalizing disorders. The American Journal of Psychiatry 163(5), pp. 857–864.

Hill, W.D., Arslan, R.C., Xia, C., et al. 2018. Genomic analysis of family data reveals additional genetic effects on intelligence and personality. Molecular Psychiatry.

KING KING: Relationship Inference Software [Online]. Available at: http://people.virginia.edu/~wc9c/KING/ [Accessed: 25 October 2018].

Knopik, V. S., Neiderhiser, J. M., DeFries, J. C., & Plomin, R. ed. 2017. Behavioral genetics (7th ed.). New York: Worth.

Laurin, C.A., Hottenga, J.-J., Willemsen, G., Boomsma, D.I. and Lubke, G.H. 2015. Genetic analyses benefit from using less heterogeneous phenotypes: an illustration with the hospital anxiety and depression scale (HADS). Genetic Epidemiology 39(4), pp. 317–324.

Lee, J.J., Wedow, R., Okbay, A., et al. 2018. Gene discovery and polygenic prediction from a genome-wide association study of educational attainment in 1.1 million individuals. Nature Genetics 50(8), pp. 1112–1121.

Lee, S.H., Weerasinghe, W.M.S.P., Wray, N.R., Goddard, M.E. and van der Werf, J.H.J. 2017. Using information of relatives in genomic prediction to apply effective stratified medicine. Scientific reports 7, p. 42091.

Lewis, C.M. and Vassos, E. 2017. Prospects for using risk scores in polygenic medicine. Genome Medicine 9(1), p. 96.

Luciano, M., Hagenaars, S.P., Davies, G., et al. 2018. Association analysis in over 329,000 individuals identifies 116 independent variants influencing neuroticism. Nature Genetics 50(1), pp. 6–11.

Mackenbach, J.P., Stirbu, I., Roskam, A.-J.R., et al. 2008. Socioeconomic inequalities in health in 22 European countries. The New England Journal of Medicine 358(23), pp. 2468–2481.

Mathieson, I. and McVean, G. 2012. Differential confounding of rare and common variants in spatially structured populations. Nature Genetics 44(3), pp. 243–246.

McAdams, T.A., Hannigan, L.J., Eilertsen, E.M., Gjerde, L.C., Ystrom, E. and Rijsdijk, F.V. 2018. Revisiting the Children-of-Twins Design: Improving Existing Models for the Exploration of Intergenerational Associations. Behavior Genetics 48(5), pp. 397–412.

Nagel, M., Jansen, P.R., Stringer, S., et al. 2018. Meta-analysis of genome-wide association studies for neuroticism in 449,484 individuals identifies novel genetic loci and pathways. Nature Genetics 50(7), pp. 920–927.

Nivard, M.G., Middeldorp, C.M., Dolan, C.V. and Boomsma, D.I. 2015. Genetic and environmental stability of neuroticism from adolescence to adulthood. Twin Research and Human Genetics 18(6), pp. 746–754.

Ormel, J., Jeronimus, B.F., Kotov, R., et al. 2013. Neuroticism and common mental disorders: meaning and utility of a complex relationship. Clinical Psychology Review 33(5), pp. 686–697.

Plomin, R., Corley, R., Caspi, A., Fulker, D.W. and DeFries, J. 1998. Adoption results for selfreported personality: evidence for nonadditive genetic effects? Journal of personality and social psychology 75(1), pp. 211–218.

Plomin, R., DeFries, J.C., Knopik, V.S. and Neiderhiser, J.M. 2016. Top 10 replicated findings from behavioral genetics. Perspectives on psychological science?: a journal of the Association for Psychological Science 11(1), pp. 3–23.

Polderman, T.J.C., Benyamin, B., de Leeuw, C.A., et al. 2015. Meta-analysis of the heritability of human traits based on fifty years of twin studies. Nature Genetics 47(7), pp. 702–709.

Robinson, M.R., Kleinman, A., Graff, M., et al. 2017. Genetic evidence of assortative mating in humans. Nature Human Behaviour 1(1), p. 0016.

Smith, B.H., Campbell, H., Blackwood, D., et al. 2006. Generation Scotland: the Scottish Family Health Study; a new resource for researching genes and heritability. BMC Medical Genetics 7, p. 74.

Smith, D.J., Escott-Price, V., Davies, G., et al. 2016. Genome-wide analysis of over 106 000 individuals identifies 9 neuroticism-associated loci. Molecular Psychiatry 21(6), pp. 749–757.

Smith, D.J., Nicholl, B.I., Cullen, B., et al. 2013. Prevalence and characteristics of probable major depression and bipolar disorder within UK biobank: cross-sectional study of 172,751 participants. Plos One 8(11), p. e75362.

van den Berg, S.M., de Moor, M.H.M., McGue, M., et al. 2014. Harmonization of Neuroticism and Extraversion phenotypes across inventories and cohorts in the Genetics of Personality Consortium: an application of Item Response Theory. Behavior Genetics 44(4), pp. 295–313.

van der Sluis, S., Verhage, M., Posthuma, D. and Dolan, C.V. 2010. Phenotypic complexity, measurement bias, and poor phenotypic resolution contribute to the missing heritability problem in genetic association studies. Plos One 5(11), p. e13929.

Visscher, P.M., Medland, S.E., Ferreira, M.A.R., et al. 2006. Assumption-free estimation of heritability from genome-wide identity-by-descent sharing between full siblings. PLoS Genetics 2(3), p. e41.

Xia, C., Amador, C., Huffman, J., et al. 2016. Pedigree-and SNP-Associated Genetics and Recent Environment are the Major Contributors to Anthropometric and Cardiometabolic Trait Variation. PLoS Genetics 12(2), p. e1005804.

Yang, J., Bakshi, A., Zhu, Z., et al. 2015. Genetic variance estimation with imputed variants finds negligible missing heritability for human height and body mass index. Nature Genetics 47(10), pp. 1114–1120.

Yang, J., Benyamin, B., McEvoy, B.P., et al. 2010. Common SNPs explain a large proportion of the heritability for human height. Nature Genetics 42(7), pp. 565–569.

Yang, J., Lee, S.H., Goddard, M.E. and Visscher, P.M. 2011. GCTA: a tool for genome-wide complex trait analysis. American Journal of Human Genetics 88(1), pp. 76–82.

Yengo, L., Robinson, M.R., Keller, M.C., et al. 2018. Imprint of assortative mating on the human genome. Nature human behaviour 2(12), pp. 948–954.

Young, A.I., Frigge, M.L., Gudbjartsson, D.F., et al. 2018. Relatedness disequilibrium regression estimates heritability without environmental bias. Nature Genetics 50(9), pp. 1304–1310.

Zaitlen, N., Kraft, P., Patterson, N., et al. 2013. Using extended genealogy to estimate components of heritability for 23 quantitative and dichotomous traits. PLoS Genetics 9(5), p. e1003520.

Zuk, O., Hechter, E., Sunyaev, S.R. and Lander, E.S. 2012. The mystery of missing heritability: Genetic interactions create phantom heritability. Proceedings of the National Academy of Sciences of the United States of America 109(4), pp. 1193–1198.

