## Supplementary Information for "Familial influences on Neuroticism and Education in the UK Biobank"

**Contents:**

Supplementary Table 1: Full GREML-KIN model-fitting results for Neuroticism. Page 2.

Supplementary Table 2: Full GREML-KIN model-fitting results for Education (I). Page 4.

Supplementary Table 3: Full GREML-KIN model-fitting results for Education (II). Page 5.

Supplementary Figure 1: GREML-KIN model-fitting results for alternative Education variables with fewer response categories. Page 7.

References. Page 8.

Supplementary Table 1: Full GREML-KIN model-fitting results for Neuroticism

Note: G (unrelated) refers to results from a standard GREML model using a relatedness cutoff of <0.025; selected model = GK (bold, underlined); GKFSC = full model; LL=log likelihood.

| Model | h2g (se) | h2kin (se) | ef2 (se) | es2 (se) | ec2 (se) | LL |  |
| --- | --- | --- | --- | --- | --- | --- | --- |
| G (unrelated) | 0.10 (0.01) |  |  |  |  | -73310.3 | 44694 |
| GKFSC | 0.11 (0.01) | 0.18 (0.08) | 0.01 (0.04) | 0.00 (0.2) | 0.04 (0.04) | -108742.05 | 66118 |
| KFSC |  | 0.25 (0.07) | 0.01 (0.05) | 0.00 (0.02) | 0.02 (0.04) | -108824.31 | 66118 |
| GFSC | 0.11 (0.01) |  | 0.08 (0.02) | 0.00 (0.02) | 0.05 (0.02) | -108744.832 | 66118 |
| GKSC | 0.11 (0.01) | 0.17 (0.03) |  | 0.00 (0.02) | 0.03 (0.01) | -108742.06 | 66118 |
| GKFC | 0.11 (0.01) | 0.18 (0.08) | 0.00 (0.04) | 0.04 (0.04) |  | -108742.053 | 66118 |
| GKFS | 0.11 (0.01) | 0.12 (0.03) | 0.03 (0.01) | 0.01 (0.02) |  | -108742.42 | 66118 |
| FSC |  |  | 0.12 (0.01) | 0.00 (0.02) | (-)0.10 (0.01) | -108827.66 | 66118 |
| KSC |  | 0.26 (0.03) |  | 0.00 (0.02) | 0.03 (0.01) | -108824.33 | 66118 |
| KFC |  | 0.25 (0.7) | 0.01 (0.04) |  | 0.02 (0.04) | -108824.32 | 66118 |
| KFS |  | 0.21 (0.03) | 0.03 (0.01) | 0.00 (0.02) |  | -108824.44 | 66118 |
| GSC | 0.12 (0.01) |  |  | 0.07 (0.01) | 0.03 (0.01) | -108758.09 | 66118 |
| GFC | 0.11 (0.01) |  | 0.08 (0.01) |  | (-)0.05 (0.01) | -108744.84 | 66118 |
| GFS | 0.11 (0.01) |  | 0.04 (0.01) | 0.04 (0.04) |  | -108749.29 | 66118 |
| GKC | 0.11 (0.01) | 0.16 (0.02) |  |  | 0.03 (0.01) | -108742.08 | 66118 |
| GKS | 0.11 (0.01) | 0.17 (0.03) |  | 0.00 (0.02) |  | -108748.17 | 66118 |
| GKF | 0.11 (0.01) | 0.11 (0.02) | 0.03 (0.01) |  |  | -108742.5 | 66118 |
| SC |  |  |  | 0.13 (0.01) | 0.03 (0.01) | -108863.81 | 66118 |
| FC |  |  | 0.13 (0.01) |  | (-)0.10 (0.01) | -108827.67 | 66118 |
| FS | Error: the variance-covariance matrix V is not positive definite. | | | | |  | 66118 |
| KC |  | 0.26 (0.01) |  |  | 0.03 (0.01) | -108824.34 | 66118 |
| KS |  | 0.26 (0.03) |  | 0.00 (0.02) |  | -108830.31 | 66118 |
| KF |  | 0.21 (0.02) | 0.03 (0.01) |  |  | -108824.44 | 66118 |
| GC | 0.15 (0.01) |  |  |  | 0.03 (0.01) | -108792.88 | 66118 |
| GS | 0.12 (0.01) |  |  | 0.07 (0.01) |  | -108764.24 | 66118 |
| GF | 0.12 (0.01) |  | 0.05 (0.01) |  |  | -108754.91 | 66118 |
| **GK** | **0.11 (0.01)** | **0.16 (0.02)** |  |  |  | **-108748.19** | **66118** |
| C |  |  |  |  | 0.03 (0.01) | -108998.069 | 66118 |
| S |  |  |  | 0.13 (0.008) |  | -108869.9 | 66118 |
| F | Error: the variance-covariance matrix V is not positive definite. | | | | |  | 66118 |
| K | Error: the variance-covariance matrix V is not positive definite. | | | | |  | 66118 |
| G | 0.15 (0.008) | |  |  |  | -108798.98 | 66118 |

Supplementary Table 2: Full GREML-KIN model-fitting results for Education (I)

Note: G (unrelated) refers to results from a standard GREML model using a relatedness cutoff of <0.025; selected model = GKC (bold, underlined); GKFSC = full model; LL=log likelihood.

| Model | h2g | h2kin | ef2 | es2 | ec2 | se (h2g) | se (h2kin) | se (ef2) | se (es2) | se (ec2) | LL | N |
| --- | --- | --- | --- | --- | --- | --- | --- | --- | --- | --- | --- | --- |
| G (unrelated) | 0.19 |  |  |  |  | 0.01 |  |  |  |  | -32911.39 | 42853 |
| GKFSC | 0.12 | 0.42 | 0.01 | 0.01 | 0.37 | 0.01 | 0.09 | 0.05 | 0.02 | 0.05 | -28928.39 | 62353 |
| KFSC | 0.55 | 0.01 | 0.01 | 0.38 |  |  | 0.09 | 0.05 | 0.02 | 0.05 | -29040.7 | 62353 |
| GFSC | 0.13 | 0.21 | 0.01 | 0.16 |  | 0.01 |  | 0.01 | 0.02 | 0.02 | -28938.363 | 62353 |
| GKSC | 0.12 | 0.43 |  | 0.01 | 0.38 | 0.01 | 0.03 |  | 0.02 | 0.01 | -28928.41 | 62353 |
| GKFC | 0.12 | 0.42 | 0.01 |  | 0.36 | 0.01 | 0.09 | 0.05 |  | 0.05 | -28928.55 | 62353 |
| GKFS | 0.12 | -0.12 | 0.36 | -0.07 |  | 0.01 | 0.01 | 0.01 | 0.01 |  | -28978.33 | 62353 |
| FSC |  |  | 0.28 | 0.01 | 0.11 |  |  | 0.01 | 0.02 | 0.02 | -29059.45 | 62353 |
| KSC |  | 0.57 |  | 0.01 | 0.39 |  | 0.03 |  | 0.02 | 0.01 | -29040.72 | 62353 |
| KFC |  | 0.55 | 0.01 |  | 0.37 |  | 0.09 | 0.05 |  | 0.05 | -29040.88 | 62353 |
| KFS |  | -0.04 | 0.37 | -0.06 |  |  | 0.01 | 0.01 | 0.01 |  | -29084.31 | 62353 |
| GSC | 0.16 |  |  | 0.21 | 0.38 | 0.01 |  |  | 0.01 | 0.01 | -29028.94 | 62353 |
| GFC | 0.13 |  | 0.22 |  | 0.16 | 0.01 |  | 0.01 |  | 0.01 | -28938.55 | 62353 |
| GFS | 0.1 |  | 0.34 | -0.1 |  | 0.01 |  | 0.01 | 0.01 |  | -29006.73 | 62353 |
| **GKC** | **0.12** | **0.45** |  |  | **0.38** | **0.01** | **0.02** |  |  | **0.01** | **-28928.6** | **62353** |
| GKS | 0.17 | 0.39 |  | 0.01 |  | 0.01 | 0.03 |  | 0.02 |  | -30148.75 | 62353 |
| GKF | 0.11 | -0.14 | 0.34 |  |  | 0.01 | 0.01 | 0.01 |  |  | -29000.26 | 62353 |
| SC |  |  |  | 0.29 | 0.39 |  |  |  | 0.01 | 0.01 | -29220.23 | 62353 |
| FC |  |  | 0.29 |  | 0.1 |  |  | 0.01 |  | 0.01 | -29059.65 | 62353 |
| FS | Error: the variance-covariance matrix V is not positive definite. | | | | |  |  |  |  |  |  |  |
| KC |  | 0.58 |  |  | 0.39 |  | 0.01 |  |  | 0.01 | -29040.92 | 62353 |
| KS |  | 0.57 |  | 0.02 |  |  | 0.03 |  | 0.02 |  | -30330.59 | 62353 |
| KF |  | -0.06 | 0.35 |  |  |  | 0.01 | 0.01 |  |  | -29099.56 | 62353 |
| GC | 0.25 |  |  |  | 0.36 |  |  |  |  |  | -29298.03 | 62353 |
| GS | 0.2 |  |  | 0.19 |  | 0.01 |  |  | 0.01 |  | -30233.13 | 62353 |
| GF | 0.08 |  | 0.31 |  |  | 0.01 |  | 0.01 |  |  | -29063.6 | 62353 |
| GK | 0.17 | 0.41 |  |  |  | 0.01 | 0.02 |  |  |  | -30149.26 | 62353 |
| C |  |  |  |  | 0.4 |  |  |  |  | 0.01 | -29855.366 | 62353 |
| S |  |  |  | 0.3 |  |  |  |  |  |  | -30516.59 | 62353 |
| F | Error: the variance-covariance matrix V is not positive definite. | | | | |  |  |  | 0.01 |  |  | 62353 |
| K |  | 0.6 |  |  |  |  | 0.01 |  |  |  | -30331.19 | 62353 |
| G | 0.29 |  |  |  |  | 0.01 |  |  |  |  | -30453.93 | 62353 |

Supplementary Table 3: Full GREML-KIN model-fitting results for Education (II)

Note: G (unrelated) refers to results from a standard GREML model using a relatedness cutoff of <0.025; selected model = GKC (bold, underlined); GKFSC = full model; LL=log likelihood.

| Model | h2g | h2kin | ef2 | es2 | ec2 | se (h2g) | se (h2kin) | se (ef2) | se (es2) | se (ec2) | LL | N |
| --- | --- | --- | --- | --- | --- | --- | --- | --- | --- | --- | --- | --- |
| G (unrelated) | 0.2 |  |  |  |  | 0.03 |  |  |  |  | -10585.4 | 21202 |
| GKFSC | 0.12 | 0.39 | 0 | 0.06 | 0.37 | 0.02 | 0.12 | 0.06 | 0.02 | 0.06 | -14326.51 | 30793 |
| KFSC |  | 0.12 | 0.42 | 0.03 | 0.33 |  | 0.02 | 0.12 | 0.06 | 0.06 | -14330.5 | 30793 |
| GFSC | 0.12 |  | 0.19 | 0.06 | 0.18 | 0.02 |  | 0.02 | 0.02 | 0.02 | -14329.998 | 30793 |
| GKSC | 0.12 | 0.39 |  | 0.06 | 0.37 | 0.02 | 0.04 |  | 0.02 | 0.01 | -14326.51 | 30793 |
| GKFC | 0.12 | 0.42 | 0.03 |  | 0.33 | 0.02 | 0.12 | 0.06 |  | 0.06 | -14330.5 | 30793 |
| GKFS | 0.12 | -0.22 | 0.35 | 0.02 |  | 0.02 | 0.03 | 0.01 | 0.02 |  | -14341.56 | 30793 |
| FSC |  |  | 0.25 | 0.06 | 0.01 |  |  | 0.02 | 0.02 | 0.02 | -14357.32 | 30793 |
| KSC |  | 0.52 |  | 0.06 | 0.37 |  | 0.04 |  | 0.02 | 0.01 | -14351.02 | 30793 |
| KFC |  | 0.53 | 0.04 |  | 0.33 |  | 0.12 | 0.06 |  | 0.06 | -14354.95 | 30793 |
| KFS |  | -0.1 | 0.35 | 0.02 |  |  | 0.02 | 0.01 | 0.02 |  | -14367.11 | 30793 |
| GSC | 0.17 |  |  | 0.23 | 0.36 | 0.02 |  |  | 0.01 | 0.01 | -14364.27 | 30793 |
| GFC | 0.12 |  | 0.02 |  | 0.13 | 0.02 |  | 0.01 |  | 0.02 | -14334.55 | 30793 |
| GFS | 0.07 |  | 0.33 | -0.04 |  | 0.02 |  | 0.01 | 0.01 |  | -14363.75 | 30793 |
| **GKC** | **0.12** | **0.49** |  |  | **0.37** | **0.02** | **0.03** |  |  | **0.01** | **-14330.63** | **30793** |
| GKS | 0.17 | 0.35 |  | 0.07 |  | 0.02 | 0.04 |  | 0.02 |  | -14844.34 | 30793 |
| GKF | 0.12 | -0.21 | 0.35 |  |  | 0.02 | 0.03 | 0.01 |  |  | -14342.22 | 30793 |
| SC |  |  |  | 0.32 | 0.37 |  |  |  | 0.01 | 0.01 | -14425.32 | 30793 |
| FC |  |  | 0.3 |  | 0.07 |  |  | 0.01 |  | 0.01 | -14362.03 | 30793 |
| FS | Error: the variance-covariance matrix V is not positive definite. | | | | |  |  |  |  |  |  | 30793 |
| KC |  | 0.37 |  |  | 0.37 |  | 0.02 |  |  | 0.01 | -14355.15 | 30793 |
| KS |  | 0.52 |  | 0.07 |  |  | 0.04 |  | 0.02 |  | -14886.21 | 30793 |
| KF |  | -0.09 | 0.36 |  |  |  | 0.02 | 0.01 |  |  | -14367.59 | 30793 |
| GC | 0.34 |  |  |  | 0.34 | 0.01 |  |  |  | 0.01 | -14505.64 | 30793 |
| GS | 0.22 |  |  | 0.21 |  | 0.01 |  |  | 0.01 |  | -14874.78 | 30793 |
| GF | 0.05 |  | 0.31 |  |  | 0.01 |  | 0.01 |  |  | -14368.47 | 30793 |
| GK | 0.17 | 0.45 |  |  |  | 0.02 | 0.03 |  |  |  | -14850.19 | 30793 |
| C |  |  |  |  | 0.38 |  |  |  |  | 0.01 | -14836.699 | 30793 |
| S |  |  |  | 0.33 |  |  |  |  | 0.01 |  | -14964.37 | 30793 |
| F | Error: the variance-covariance matrix V is not positive definite. | | | | |  |  |  |  |  |  | 30793 |
| K |  | 0.63 |  |  |  |  | 0.02 |  |  |  | -14892.16 | 30793 |
| G | 0.39 |  |  |  |  | 0.01 |  |  |  |  | -14994.05 | 30793 |

Supplementary Figure 1: GREML-KIN model-fitting results for alternative education variables with fewer response categories.

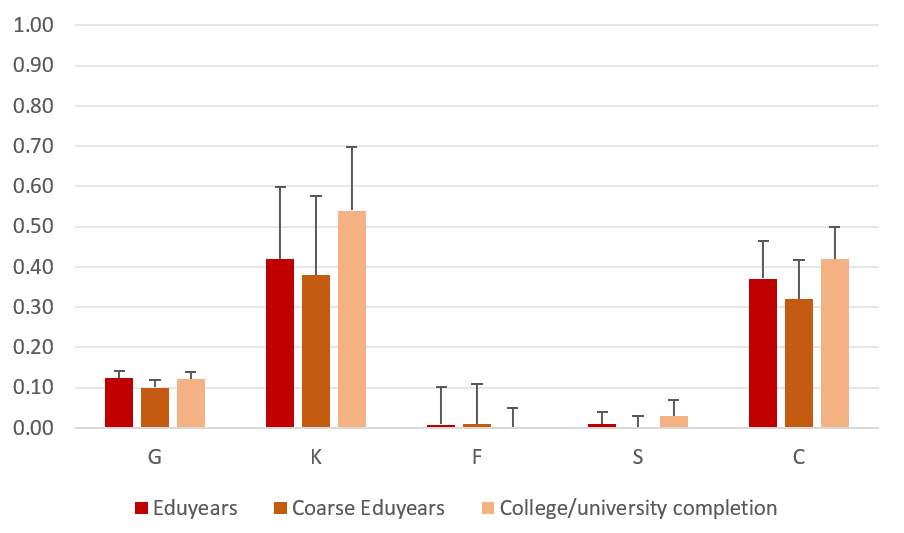

Note: full models with all five variance components are presented here. As indicated in the main text, G= population-level effects of common genotyped SNPs; K= kin-based genetic effects; F= nuclear family (siblings, parent-offspring, couple); S= sibling similarity; C= couple similarity. The coarse eduyears variable was constructed as in the Lee et al. (2018) paper, by collapsing the full eduyears variable from 6 to 3 categories (representing 10, 13 and 19 years of education, respectively). The college/university completion variable was constructed as a binary variable, with individuals scoring 1 if they report university as their highest level of qualification, and 0 if their highest qualification was below that.

Supplementary Figure 1 shows that estimates for the 3 phenotypes are not significantly different -- they have overlapping 95% confidence intervals. However, it is interesting that that heritability is higher for the binary college completion variable (66%) than for more detailed and accurate phenotypes capturing more variation. Heritabilities were 54% and 48% for years of education measured with 6 and 3 categories, respectively. Supplementary analyses in Lee et al. (2018) indicated that heritability was significantly reduced in UK Biobank when response options were dropped. The inconsistency with our results could be explained by their use of the full sample, rather than a much smaller sample of relatives, or by their use of a different heritability method (LD score regression).

References

Lee, J.J., Wedow, R., Okbay, A., et al. 2018. Gene discovery and polygenic prediction from a genome-wide association study of educational attainment in 1.1 million individuals. *Nature Genetics* 50(8), pp. 1112–1121.
